# Enhanced anti-tumor activity by Zinc Finger Repressor-driven epigenetic silencing of immune checkpoints and TGFBR2 in CAR-T cells and TILs

**DOI:** 10.1101/2024.10.11.613893

**Authors:** Marion David, Phillip Schiele, Davide Monteferrario, Gaëlle Saviane, Angélique E. Martelli, Coralie F. Dupont, Caroline Jeanneau, Irène Marchetti, Satish K. Tadi, Julia Vahldick, Lynn N. Truong, Yuanyue Zhou, Igor M. Sauer, Wenzel Schöning, Il-Kang Na, Andreas Reik, Marco Frentsch, Maurus de la Rosa, David Fenard

**Author notes:** **Correspondence:** David Fenard, PhD, Sangamo Therapeutics France, Les Cardoulines HT1, Allée de la Nertière, 06560 Valbonne France, **Email:** or. Co-First authors (Marion David, Phillip Schiele, Davide Monteferrario). Co-Senior authors (Marco Frentsch, Maurus de la Rosa, David Fenard).

## Abstract

CAR-T therapies have shown remarkable success in treating hematological malignancies. However, effectiveness against solid tumors remains limited due to the immunosuppressive tumor microenvironment (TME), such as TGF-β signaling and upregulated immune checkpoints (ICs). Furthermore, identifying universal, tumor-specific targets for CAR-T cells in solid tumors is challenging, but using reinvigorated, immunosuppressive-resistant tumor-infiltrating lymphocytes (TILs) could be a promising alternative approach. Unlike nucleases, which may induce genotoxic DNA double-strand breaks, multiplexed Zinc Finger Repressors (ZFR) offer a safer alternative for knocking out TME-related immunosuppressive factors. We epigenetically repressed PD-1 expression both in CAR-T cells and TILs from colorectal liver metastases. PD-1 repression did not affect T cell viability, proliferation, or functionality. In a murine B cell lymphoma model, PD-1-repressed CD19-CAR-T cells exhibited enhanced anti-tumor activity and improved survival. Notably, PD-1 repression alone did not increase cytotoxicity against a PD-L1-positive colorectal cell line in vitro. To further increase anti-tumor potency in this context, ZFR-expressing lentiviral vectors targeting PD-1 and other ICs (LAG-3, TIM-3, TIGIT) or TGFBR2 were developed, improving significantly the cytotoxic activity in TILs. This strategy highlights the potential to enhance tumor-reactive T cells and improve anti-cancer immunotherapies by epigenetically repressing immunosuppressive factors in the TME using multiplexed ZFRs.

## INTRODUCTION

Chimeric antigen receptor T (CAR-T) cell therapies have revolutionized the treatment of leukemia and lymphoma by delivering a defined dose of T cells engineered for targeting a specific surface antigen.^1^ However, this approach shows limited efficacy in solid tumors.^2^ Besides the selection of suitable cancer tissue-specific antigens that are expressed in different patient cohorts and different tumor stages in solid tumors, the immunosuppressive mechanisms mediated by the tumor microenvironment (TME) is a major limitation for CAR-T cell therapies. T cells capable of migrating into tumor tissue are exposed to immune suppressive mediators of the TME like TGF-β, which inhibit their effector functions.^3^ Moreover, upon recognition of their cognate antigen and repeated stimulation by tumor antigens, T cells undergo transcriptional and epigenetic changes that lead to an upregulation of inhibitory receptors, also called immune checkpoints (ICs), such as PD-1, TIM-3, LAG-3 and TIGIT. The ligands of ICs expressed by tumor-resident cells bind to ICs, leading to the silencing of immune activation and of the anti-cancer cytotoxicity of tumor-specific T cells.^4, 5^ Accordingly, tumor-infiltrating lymphocytes (TILs), which are enriched with T cell receptor (TCR) clones specific to tumor antigens, often exhibit an exhausted or dysfunctional phenotype, similar to that observed in the limitations of CAR-T therapy.^6–8^ Hence, targeting ICs with monoclonal antibodies has transformed cancer treatment by enhancing the immune system’s ability to eliminate cancer cells.^9, 10^ Despite their remarkable efficacy in various cancers, not all patient populations respond to systemic IC blockade, and many patients experience acute or chronic side effects related to the treatment.^11^ Additionally, due to the pleiotropic and context-dependent roles of soluble factors such as TGFβ in tissue homeostasis and tumor prevention, therapeutic strategies targeting these factors may benefit from the specific genetic engineering of anti-tumor immune cells. Targeted manipulation of multiple immunosuppressive pathways in tumor-reactive T cells, such as CAR-T cells or TILs, is therefore a promising approach to overcome unresponsiveness and immune-induced toxicities of current systemic therapies.

Previously, removal of PD-1 alone or in combination with other ICs using small hairpin RNA (shRNA) has been shown to enhance the anti-tumor capacity of CAR-T cells.^12, 13^ However, the continuous expression of shRNAs in CAR-T cells can potentially affect their persistence. This toxicity arises from off-target effects, interference with, and saturation of the endogenous micro-RNA (miRNA) pathway, as well as the activation of the interferon response.^14^ Additionally, incorporating multiple shRNAs into a lentiviral backbone is challenging due to sequence homologies, which are known to induce lentiviral vector (LV) recombination.^15^ Today, targeted DNA endonucleases are the most widely used gene editing technology to knock-out gene expression. It consists of a nuclease domain fused to a DNA-binding domain (DBD) derived either from a Zinc-Finger array (ZFN),^16^ a TALE domain (TALEN)^17^ or derived from the RNA-guided CRISPR-Cas system.^18^ Although these technologies provide efficient gene knockout with high specificity, the introduction of double-strand breaks into the genome may carry the risk of mutational events and chromosomal rearrangements.^19–21^ These mutations increase the likelihood of genotoxic side effects, a risk further exacerbated by the simultaneous introduction of multiple edits.

An attractive alternative is to fuse DBDs to an epigenetic repressor instead of nuclease, such as members of the Krüppel-associated box (KRAB).^22^ Epigenetic gene regulation involves reversible, heritable changes in gene expression that occur without altering the genomic DNA sequence, including DNA methylation and histone modifications, modulating chromatin structure and accessibility to regulate gene transcription and cellular functions. The KRAB epigenetic repressors can be fused either to a RNA-guided dead Cas9 (CRISPRi),^23^ a TALE DBD (TALE-KRAB),^24^ or a Zinc-Finger protein, also known as ZF-Repressor (ZFR).^25–28^ The last generation of ZFRs possess several advantages including a modular and compact design allowing multiplexing in LV backbone,^26^ a versatile DNA binding capacity,^29^ and a human and not bacterial origin reducing the chance of pre-existing immunity.^30^

To address TME challenges, we developed an approach to enhance CAR-T and TIL therapies by epigenetically silencing IC pathways, shielding tumor-reactive cells from the TME. Applied to TILs, this method leverages their natural polyclonal specificity, bypassing the need for tumor-specific CAR selection. Thus, targeted manipulation of multiple immunosuppressive pathways in tumor-reactive T cells presents a promising strategy to overcome the limitations of current systemic therapies. Here, we describe the generation of multiplexed ZFR-lentiviral vectors for the downregulation of PD-1 alone or in combination with LAG-3, TIM-3, TIGIT, and TGFBR2 to improve the efficacy of CD19-CAR-T cells and TILs isolated from colorectal cancer liver metastases (CRLMs) patients.

## RESULTS

### Stable phenotype and functionality of CAR-T cells after ZFR-mediated epigenetic silencing of PD-1

To evaluate the impact of the PD-1 epigenetic repression in CAR-T cells, an extensive screening of more than hundred ZFR candidates targeting the transcription start site (TSS) region of the *PDCD1* gene has been performed, allowing the identification of a lead candidate (see the star in Figure S1 and Supplemental Material and Methods). Next, human T cells were transduced with a lentiviral construct expressing the CD19-CAR alone or in combination with the selected PD-1 ZFR (Figure 1A and Figure S2). As measured by flow cytometry, PD-1 repression in PD-1 ZFR-expressing T cells was potent (Figure 1B) and stable over time (Figure 1C) compared to the control condition. To further confirm the effectiveness of the PD-1 ZFR, PD-1 expression was induced either through a TCR stimulation (anti-CD3/CD28 beads) or a CD19-CAR stimulation, after incubation with Nalm6-L-PD-L1, a CD19-positive cell line engineered to express PD-L1, one of the natural PD-1 ligands (Figure S3). Regardless of the type of stimulation, cell surface PD-1 expression was strongly repressed in PD-1 ZFR-expressing T cells (Figure 1D). To evaluate the impact of the PD-1 ZFR expression on CAR-T cell functionality, we first compared the efficiency of CD19-CAR T cells to kill NALM6-L-PD-L1 cancer cells *in vitro* in absence or presence of PD-1 ZFR. As shown in Figure 1E, the cytotoxic activity of PD-1 ZFR-expressing CAR-T cells was comparable to the control condition, even at low effector-to-target ratios (E:T = 1:32). Next, the activation characteristics of PD-1 ZFR-expressing CAR-T cells were evaluated 24-hrs after incubation with Nalm6-L-PD-L1 tumor cells. We showed that PD-1 ZFR expression did not modulate the capacity to express the CD69 activation marker, the CD107a degranulation marker (Figure 1F) or the production of pro-inflammatory cytokines, namely IFNγ, IL-2, and TNFα (Figure 1G). These results indicate that the PD-1 epigenetic repression did not alter the phenotype and functionality of CAR-T cells *in vitro*.

**Figure 1.**
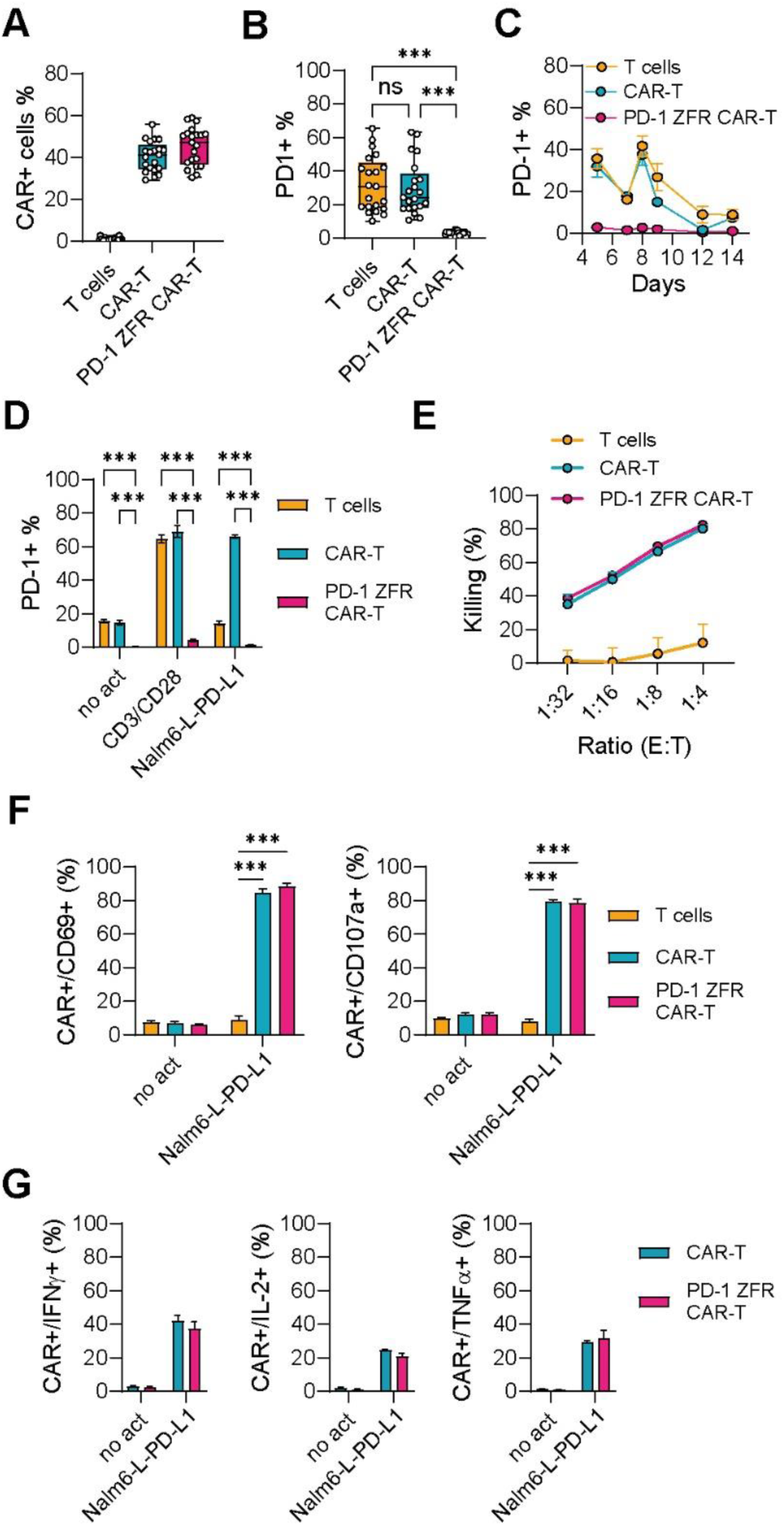
T cell phenotype and functionality after ZFR-mediated repression of PD-1. Human T cells were transduced with LV constructs expressing the CD19-CAR alone or in combination with the PD-1 ZFR. CAR expression (A) and PD-1 repression efficiency (B) were monitored in T cells using flow cytometry (D3 post-LV) Two-way ANOVA Friedman test (***p<0.001). C) Kinetic of PD-1 repression efficiency in CAR-T cells. Cells were cultured for 14 days with an anti-CD3/CD28 reactivation step at D7. D) Repression efficiency of PD-1 following TCR activation (anti-CD3/CD28 beads) or CD19-CAR activation after co-culture with a CD19-expressing cell line (Nalm6-L-PD-L1) Two-way ANOVA Tukey test (***p<0.001). E) Cytotoxic capacity of T cells, CAR-T and PD-1 ZFR-CAR-T cells after overnight co-culture with NALM-6-L-PD-L1 cells at the indicated effector: target (E:T) ratios. Results were obtained from three independent donors. Error bars indicate mean ± SD. F) Activation (left panel) and degranulation (right panel) of T cells, CAR-T and PD-1 ZFR-CAR-T cells were assessed by measuring CD69 and CD107a expression respectively, following 24hrs co-colture with NALM-6-L-PD-L1 cells at 1:3 E:T ratio. Two-way ANOVA Sidak test (***p<0.001) G) Expression of IFNγ (left panel), IL-2 (middle panel) and TNFα (right panel) in CAR-T and PD-1 ZFR-CAR-T cells was assessed by flow cytometry following a 24hrs co-colture with NALM-6-L-PD-L1 cells at 1:3 E:T ratio. Results were obtained from three independent donors. Data are represented as mean ± SEM.

### PD-1 epigenetic silencing in TILs did not alter their phenotype or polyclonal T cell functionality

TILs possess the properties of expressing an enriched repertoire of TCR clones specific to tumor antigens and the ability to migrate to and reside within tumors; however, they often exhibit exhausted or dysfunctional phenotypes due to the TME.^6, 8^ To enhance TILs functionality after transduction with a PD-1 ZFR repressor, T cells from tumor biopsies of CRLM patients were isolated (Figure 2A). Most colorectal cancers, especially CRLMs, are of the T cell excluded phenotype. Hence, TILs were isolated from the tumor cores and the closest part of the invasive margin, in line with the literature.^31, 32^ On average, 1.2 ×10^5^ (± 8×10^4^) CD3+ TILs per cm³ tumor tissue were obtained. Subsequently, TILs were activated and efficiently transduced with control and PD-1 ZFR-expressing LV with mean efficacies of 38% and 35% (data not shown). Conducting lentiviral transduction within the initial week of TILs isolation was essential for optimal efficacy. This approach minimized the starting material quantity and necessitated an expansion period before any functional characterization (Figure 2A). While PD-1 expression varied between TILs from different patients (13-67%) and slightly decreased during expansion, transduction of PD-1 ZFR resulted in 90% PD-1 repression, with the same manner after one or two weeks (Figure 2B). To assess anti-cancer reactivity of PD-1-repressed TILs, antigen-specific T cell responses were mimicked by co-culturing TILs with and EpCAM+/PD-L1+ colorectal cancer cell line (SNU-C5-GFP-Luc, see Table S3) and the addition of a bispecific anti-CD3E-EpCAM T cell engager. As shown in Figure 2C, at an E:T ratio of 1:2, TILs were effectively activated and induced SNU-C5-GFP cell lysis (Figure 2C). While a slight upward trend was observed, the PD-1 ZFR did not significantly improve cancer cell killing, indicating that PD-1 repression is not sufficient to enhance anti-cancer cytotoxicity of fully activated TILs in vitro.

**Figure 2.**
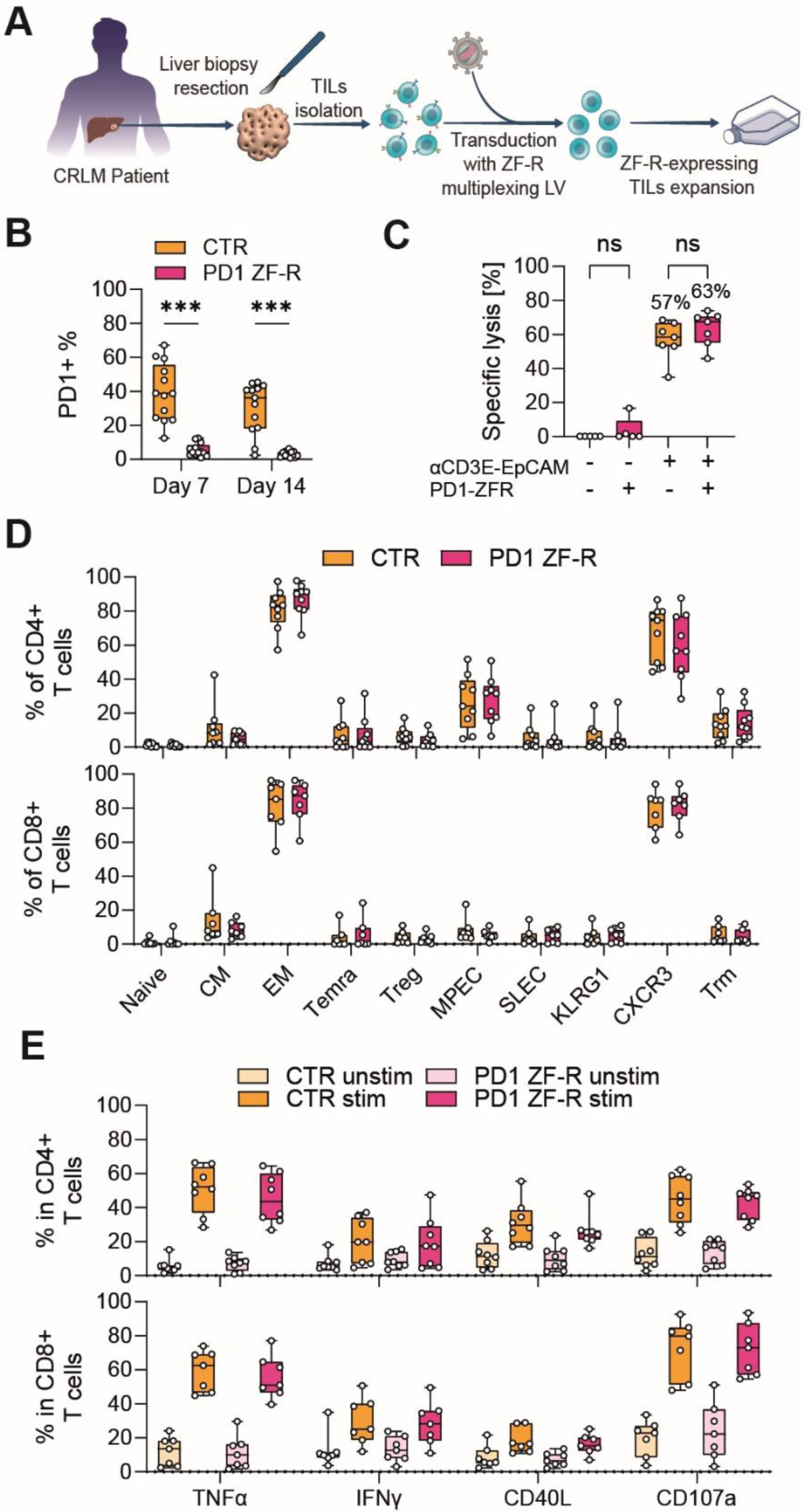
Efficient and durable PD-1 repression in TILs from CRLM patients. A) Schematic representation of the isolation, transduction and expansion of TILs isolated from CRLM patients. After tissue dissection, enriched CD3^+^ TILs were activated, transduced and expanded for three weeks. Transduction efficacy was determined seven days and functionality 21 days after transduction. B) PD-1 expression on dNGFR-enriched TILs (n=13) at day 7 and day 14 post-LV. Two-way ANOVA Sidak test (***p<0.001). C) Killing of SNU-C5-GFP-Luc cells by control or PD-1 ZFR transduced TILs in absence or presence of anti-CD3E-EpCAM antibody. Specific cancer cell line lysis was defined as reduction in viable GFP^+^ cancer cells after 48 hrs co-culture with TILs (n=8). After three weeks of culture, the phenotype (D, n = 9 donors) and the functionality (E, n = 8 donors) of expanded NGFR+ CD4+ and CD8+ TILs expressing or not the PD-1 ZFR were assessed by flow cytometry. Functionality was determined by monitoring TNFα, IFNγ, CD40L and CD107a expression before and after stimulation with anti-CD3/CD28 dynabeads for 24 hrs. Abbreviations: CM central memory T cell, EM effector memory T cell, Temra EM CD45RA+ T cell, MPEC memory precursor effector cells, SLEC short-lived effector cells, Trm Tissue-resident memory T cells. Data are represented as mean ± SEM.

Besides warranted improvements in efficacy, altering immunosuppressive molecules of T cells also bears the potential to induce an over-activated state of T cells. Therefore, we analyzed the phenotypic and functional characteristics of TILs transduced with control vector or PD-1 ZFR. As summarized in Figure 2D-E, PD-1 repressed CD4+ and CD8+ T cells did not exhibit changes in their sub-phenotypes or in the expression levels of TNFα, IFNγ, the co-stimulatory factor CD40L, and the degranulation marker CD107a, either at baseline or following reactivation with anti-CD3/CD28 beads. This indicates that PD-1 repression was effective without impacting other pathways.

### The epigenetic repression of PD-1 potentiates the anti-tumor activity of CD19 CAR-T cells in vivo

As shown in Figure 1, our experiments performed *in vitro* were unable to show a positive impact of the PD-1 epigenetic silencing on CAR-T cell functionality. This could be the consequence of a too simplistic setting, unable to recapitulate the immune interactions and TME that can be encountered *in vivo*. Therefore, to better evaluate the anti-tumor effect of CD19 CAR-T cells in presence of the PD-1 ZFR, a B-ALL xenograft mouse model was implemented. NXG immunodeficient mice were engrafted with NALM6-L-PD-L1 tumor cells, followed by CAR-T cell injection four days later (Figure 3A). Before injection, PD-1 ZFR-expressing CAR-T cells were enriched based on CAR expression (>95%) and expanded for 8 days *in vitro*. In these cells, PD-1 was almost completely repressed, as depicted in Figure S4. Bioluminescence imaging (BLI) of the mice was performed weekly to assess the tumor burden (Figure 3B). Mice injected with untransduced T cells exhibited rapid disease progression comparable to that of the saline group as showed by the BLI signal quantification, bodyweight loss, clinical score and survival curve (Figure 3C-F). In contrast, mice receiving CAR-T cells demonstrated substantial protection, resulting in a prolonged median survival (Figure 3F). Interestingly, superior tumor control was observed for CAR-T cells co-expressing PD-1 ZFR. The anti-tumor effect was maintained during the entire monitoring period, with some mice harboring a BLI signal below the threshold of detection (2/5 mice), demonstrating long-term remission (Figure 3C-F). At the time of sacrifice, the phenotype of CAR-T cells and PD-1-ZFR CAR T cells in B-ALL xenograft NXG mice has been monitored (Figure 3G). Remarkably, after six months *in vivo*, PD-1 was still highly repressed in PD-1 ZFR CAR-T cells isolated from blood (Figure 3H), spleen (Figure 3I), and bone marrow (Figure 3J), to an extent comparable to PD-1 repression observed before injection (Figure S4). These results highlight the potency and the stability of this ZFR-mediated epigenetic silencing and its strong potential in promoting anti-tumor activity when targeting PD-1.

**Figure 3.**
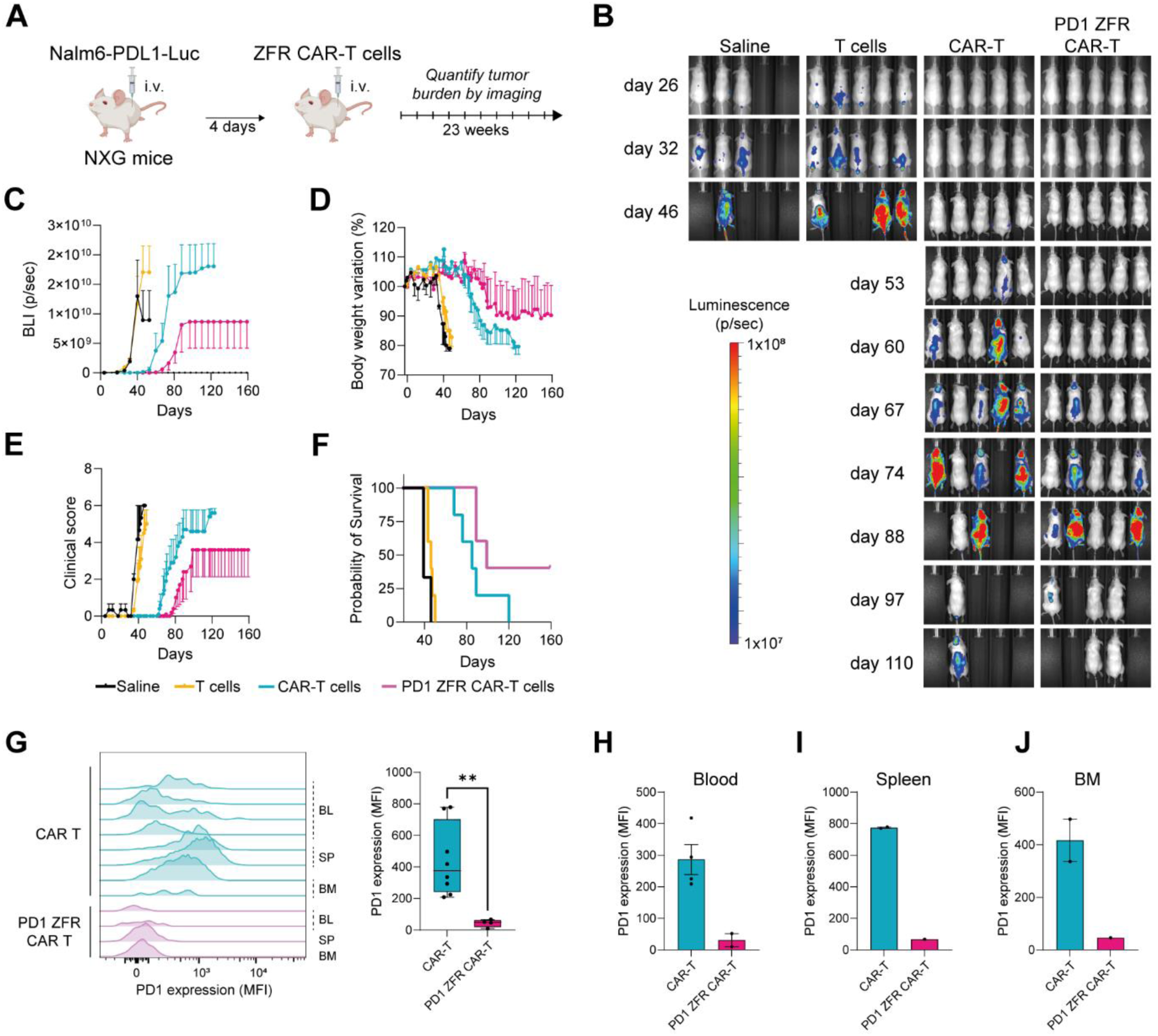
PD-1 downregulation enhances the anti-tumor activity of CAR-T cells in vivo. A) Schematic of the in vivo experimental protocol (BioRender). During six months following CAR-T or PD-1 ZFR-CAR-T cells injection, Nalm6-L-PD-L1 tumor cell proliferation was evaluated by monitoring the in vivo bioluminescence imaging (BLI) signal (photons/sec). (B) Representative mice images showing BLI from tumor cells at different timepoints in the different groups (n=3-5/group). The total BLI signal of the tumor (C), the body weight loss (D), the clinical score progression (E) and the survival rate (F) were monitored in the different mice groups over time. G) PD-1 expression levels (MFI) in CAR-T versus PD-1 ZFR-CAR-T cells in organs (BL, Blood; SP, Spleen, BM, Bone marrow) of individual mice (left) or pulled altogether (right) or by organs in blood (H), spleen (I) and bone marrow (J). Data are represented as mean ± SEM and Two-tailed Mann-Whitney test (**p<0.01).

### Multiplexed ZFR-mediated repression of ICs or TGFBR2 promotes anti-tumor cytotoxicity of TILs

In addition to PD-1, TILs from solid cancers, including colorectal cancer, express high levels of different inhibitory IC such as TIM-3, LAG-3, and TIGIT.^33–36^ Furthermore, soluble factors from the TME such as TGFβ are known to impede T cell infiltration resulting in an immune excluded tumor phenotype.^37^ Hence, we aimed to generate TILs that are resistant to the immunosuppressive TME by co-targeting diverse ICs and TGFBR2 with specific ZFRs (Figure S1). Using our established bidirectional LV design,^26^ we targeted PD-1 alone or multiplexed with TIM-3, LAG-3, TGFBR2 or TIGIT (Figure S2). The mean of transduction efficiency for multiplexed ZFR-expressing LVs ranged from 15 to 20% (Figure 4A), and importantly did not impede TILs proliferation over the course of the 21 days culture (Figure 4B). As reported previously for T cells isolated from solid tumors,^34^ TILs expressed high levels of PD-1 (Figure 4C), TIM-3, LAG-3, TIGIT and TGFBR2 (Figure 4D). Expression of PD-1 ZFR reduced PD-1 expression by 83-89% using single but also multiplexed ZFR-expressing LVs (Figure 4C). Additionally, respective multiplexed constructs effectively repressed the expression of TIM-3 (∼95%), LAG-3 (∼88%), TIGIT (∼89%) and TGFBR2 (∼74%) (Figure 4D). Importantly, suppression of multiple ICs did not result in enhanced cytokine expression after polyclonal stimulation or any change in TILs phenotypes, as observed for ZFR-expressing T cells in Figure 1 (data not shown). These data suggest that ZFR can durably suppress multiple ICs, while the T cell phenotypes and functionality of T cells remained comparable to non-modified TILs preparations. The cytotoxic activity of engineered TILs has been evaluated like in Figure 2C, after co-culture with the SNU-C5-GFP-Luc cell line in presence of anti-CD3E-EpCAM antibody. As shown in Table S3, this modified SNU-C5 cancer cell line is able to express EpCAM and more importantly, the ligands required to interact with PD-1 (PD-L1), TIM-3 (CEACAM1), LAG-3 (MHC-II, Galectin-3), TIGIT (CD112, CD155), and TGFBR2 (TGFβ variants). As shown in Figure 4E, concurrent targeting of PD-1 together with LAG-3 or TGFBR2, but not TIM-3 or TIGIT, significantly reinforced the killing efficacy compared to control or single PD-1 ZFR conditions. These results demonstrate that the multiplexing and co-delivery in TILs of ZFRs directed against ICs using a single lentiviral particle is feasible. And more importantly, the epigenetic silencing of ICs in TILs, especially the combination PD-1/LAG-3 or PD-1/TGFBR2, can promote their anti-tumor activity.

**Figure 4.**
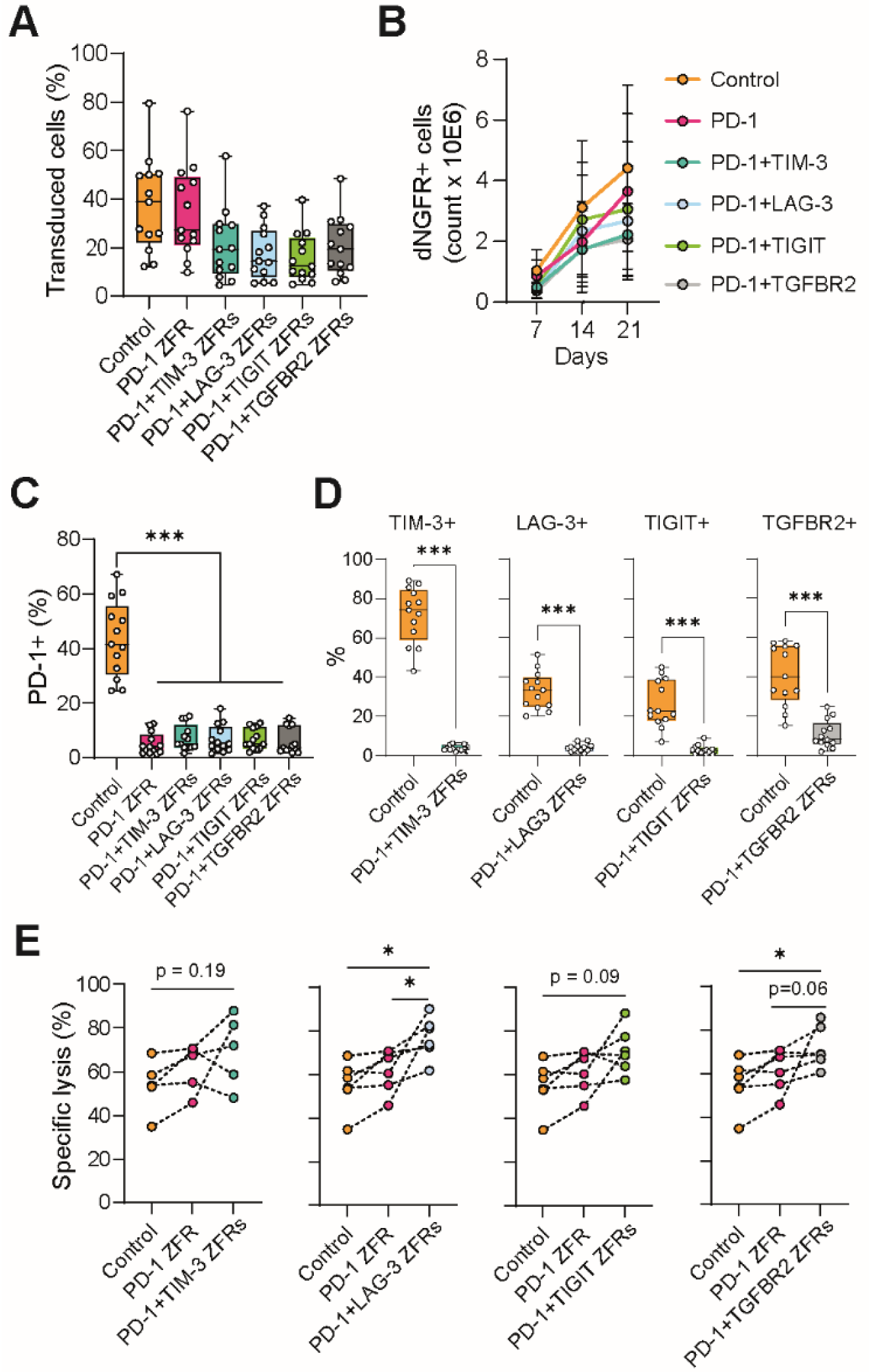
Multiplexed ICs and TGBR2 repression improve anti-cancer TILs cytotoxicity. A) Transduction efficiency of single and multiplexed ZFR-expressing LVs detected by flow cytometry (dNGFR+ cells) 7 days post-LV. B) Expansion of NGFR-enriched TILs after transduction with single PD-1 ZFR or multiplexed ZFRs. C-D) Expression of PD-1, TIM-3, LAG-3, TIGIT or TGFBR2 on dNGFR+ TILs after transduction with single PD-1 ZFR or respective multiplexed ZFRs (n=14 donors). Two-way ANOVA Friedman test (***p<0.001). E) Anti-CD3E-EpCAM-mediated killing of SNU-C5-GFP-Luc cells by TILs (n = 6-8 donors) transduced with single PD-1 ZFR or multiplexed ZFRs. Specific cancer killing was defined by reduction of viable GFP+ cancer cells compared with cell count obtained in unstimulated co-cultures in the control condition. Wilcoxon test (*p<0.05). Data are represented as mean ± SEM.

## DISCUSSION

Despite the revolutionary success of cancer immunotherapy, significant gaps remain, demanding innovative treatment strategies for certain patient populations. Cell therapy stands out for its ability to target and persist in tumors. However, a key challenge for CAR-T cell therapies is the absence of a universal tumor-specific antigen consistently expressed across patients. An alternative approach harnesses the tumor-specific TCR repertoire of autologous TILs, showing promise in melanoma after in vitro expansion.^38^ Yet, in other solid tumors such as colorectal and lung cancer, the TME poses a significant barrier to the success of both strategies. Therefore, this study investigates the application of ZFR epigenetic silencers to modulate the expression of immune checkpoints, particularly PD-1, in tumor-reactive T cells, with the aim of augmenting their anti-tumor efficacy. Following the screening of numerous ZFR candidates targeting the TSS region of the *PDCD1* gene, one lead ZFR PD-1 candidate was identified. Its expression in CAR-T led to a robust repression of cell-surface PD-1 expression, both in resting and activated T cells. The PD-1 ZFR expression did not compromise the viability, expansion capacity, TCR functionality, cytokine secretion levels, or cytotoxic activity of CAR-T cells *in vitro*. Interestingly, upon injection of CD19 CAR-T cells into a humanized B cell tumor model, a significant improvement in tumor regression and survival rates was observed in mice receiving PD-1-repressed CAR-T cells. This discrepancy with *in vitro* killing data suggests that the tumor microenvironment *in vivo* plays a critical role in modulating T cell responses. Several factors, including immunosuppressive cytokines and metabolic constraints, can attenuate the anti-tumor activity of CAR-T cells and promote their exhaustion.

This study also investigated the use of the ZFR epigenetic repressors in TILs. Lifileucel was recently approved as the first TILs therapy for solid tumors highlighting their potential to cure patients with metastatic cancers.^38^ Due to their complex nature and the challenges within the TME, optimizing those specialized T cells is key for therapeutic efficacy. Several studies indicate that improving expansion conditions, enhancing the homing or cytotoxic potential or reducing T cell exhaustion play pivotal roles in TILs products.^39, 40^ In addition to PD-1, TILs from solid cancers, especially in the advance state like metastatic colorectal cancer, express high levels of different inhibitory ICs^34^ such as TIM-3, LAG-3, and TIGIT and soluble factors from the TME such as TGFβ are known to impede T cell infiltration resulting in an immune excluded tumor phenotype.^37^ Hence, we aimed to generate TILs that are resistant to the immunosuppressive TME by lentiviral transduction of multiplexed ZFRs. Considering the plethora of inhibitory molecules within the TME, targeting of multiple pathways is a promising approach, but requires a multiplexed, specific and targeted technology like epigenetic editing. Crucially, besides the specifically targeted ICs, we did not observe any changes in T cell expansion, phenotypic markers or T cell functionality after polyclonal stimulation suggesting the absence of an aberrantly altered T cell state and an acceptable safety profile of ZFR-transduced TILs. Developing in vivo models recapitulating individualized and gene-modified IC interactions would be challenging due to the high heterogeneity of cancers and the limited amount of patient material. We therefore mimicked a multifactorial immunosuppressive TME by developing an in vitro cytotoxicity assay based on SNU-C5-GFP-Luc cells which express different IC ligands and can secrete TGFβ. In this model system, PD-1 repression alone was not sufficient to improve anti-cancer cytotoxicity. Therefore, the experimental design of the killing assay was expanded beyond PD-1 repression alone to investigate the simultaneous repression of multiple immune checkpoints, namely TIM-3, LAG-3, and TIGIT in combination with PD-1 repression. This multiplexing strategy demonstrated superior anti-tumor efficacy in vitro for PD-1 co-repressed with LAG-3, but not with TIM-3 or TIGIT. Very interestingly, it has been shown recently that the combination of LAG-3 and PD-1 inhibitors in cancer patients was able to modulate the differentiation of CD8+ T cells by enhancing responses to T cell receptor and IFNγ signaling, leading to enhanced effector functions despite the retention of an exhaustion profile.^41^ Furthermore, CD8+ T cells deficient in both PD-1 and LAG-3 enhanced anti-tumor immunity driven by an autocrine IFNγ-dependent mechanism.^42^ These observations highlight the synergistic effects of targeting multiple immune checkpoints to overcome tumor-induced immunosuppression and suggest that the targeted silencing of PD-1 and LAG-3 using ZFR epigenetic repressors is a very promising clinical approach either for TILs or CAR-T cell therapies. Concerning the inefficient data obtained with ZFR TIM-3 and TIGIT, these differences between targeted IC combinations might be attributed to their distinct functions in immune regulation and context-dependent ligand interactions.^43, 44^

This study also addressed the role of TGFβ signaling in shaping the immunosuppressive TME. By targeting TGFBR2 along with PD-1, we aimed to mitigate the inhibitory effects of TGFβ on TILs. This combinatorial approach also promoted the anti-tumor activity of TILs, which might be explained by the negative impact of TGFβ on activation.^45, 46^ Aside from directly affecting anti-cancer cytotoxicity, depletion of TGFβ signaling in tumor-reactive T cells has also the potential to shape an anti-tumorigenic TME. Dodagatta-Marri et al. identified that PD-1 blockade therapy elevates Treg levels, limiting anti-tumor responses that could be rescued by TGFβ blockade.^47^ Additionally, TGFβ is a key factor suppressing trafficking to and invasion into tumors and thereby limiting the efficacy of current immunotherapies.^37, 48^ Together, combining repression of ICs with factors rendering the TME more immunogenic has a great potential to improve the efficacy of T cell therapies. In the future, a deeper characterization of TILs biology in relation to the TME for each patient could be a valuable approach for guiding the personalized selection of the most effective IC target combinations. Altogether, these results emphasize the value of multiplexed repression of ICs by ZFR to improve the clinical efficacy of cellular anti-cancer therapies.

## MATERIAL AND METHODS

### Human T cell and TILs culture

Primary T cells were isolated from buffy coats of healthy donors (EFS, Marseille, France). Peripheral blood mononuclear cells (PBMCs) were isolated by Ficoll gradient centrifugation. T cells were isolated using the EasySep Human T cell isolation kit (StemCell Tech., Saint-Egrève, France, activated with anti-CD3/CD28 dynabeads and cultured in X-Vivo15 medium supplemented with 100 IU/ml IL-2 (Thermo Fisher Scientific, Waltham, 319 MA) and 5% Human AB Serum (Valley Biomedical, Winchester, VA).

For TILs isolation, cancerous tissues resected from patients with colorectal liver metastases (CRLM, see table S1) was approved by the Ethics Committee of Charité – Universitätsmedizin Berlin (EA1/077/22). For TILs isolation, cancerous tissues obtained from specimen after informed consent were dissociated using the Tumor Dissociation Kit together with the gentleMACS™ Dissociator (Miltenyi Biotec, Bergisch Gladbach, Germany) according to the manufacturers protocol for soft tissues. TILs were isolated with the EasySep Human CD3 Positive Selection Kit II (StemCell Tech.). Following initial activation with anti-CD3/CD28 dynabeads (Thermo Fisher Sci.), TILs were either transduced and/or expanded in complete X-Vivo15 medium supplemented with 50 ng/ml IL-15 and 10 ng/ml IL-7 (Miltenyi Biotec).^49^

### Lentiviral transduction and transduced cell enrichment

Lentiviral constructs used in this study are described in Figure S2. Third-generation SIN lentiviral vectors were produced using the classical 4-plasmid lentiviral system as described previously^26^. Lentiviral transduction was performed two days after T cell isolation and activation. Briefly, cell suspension (2×10^6^ cells/ml) were incubated with LV supernatants at a final concentration of 4×10^7^ TU/ml in complete X-Vivo15 medium. After 6 hours at 37°C and 5% CO2, cell suspensions were diluted with fresh complete medium. For TILs, transduction was repeated on two consecutive days to maximize transduction efficacy. After 3 to 5 days, transduction efficiencies were measured using flow cytometry to determine the percentage of dNGFR or CAR positive cells. For dNGFR enrichment, transduced cells were isolated with the EasySep Human CD271 Positive Selection Kit II with EasyEights EasySep Magnets (StemCell Tech.) according to the manufacturer protocol. In the case of transduced TILs, dNGFR purification was repeated after 14 days of expansion and prior to comparison of TILs functionalities. For HA-CD19 CAR-T cell enrichment, transduced cells were first labeled with an anti-HA-biotin antibody (Miltenyi Biotec) enriched using the EasySep Biotin positive selection Kit II with EasyEights EasySep Magnets (StemCell Tech.) according to the manufacturer protocol.

### *In vivo* monitoring of anti-tumor activity of ZFR-expressing CAR-T cells in mouse model

NOD-Prkdcscid-IL2rgTm1/Rj (NXG) mice were obtained from Janvier Labs (Le Genest-Saint-Isle, France). 8-week-old mice were housed in containment isolators and habituated for one week prior to experimental use. Mice were injected intravenously with 1×10^5^ Nalm6-L-PD-L1 cells expressing firefly luciferase to facilitate non-invasive tumor burden detection in vivo by bioluminescence imaging. Four days after tumor cell inoculation, 5×10^6^ T cells or HA-enriched CD19 CAR-T or PD-1 ZFR CD19 CAR-T cells were administrated intravenously. The tumor burden was monitored for one week by in vivo bioluminescence imaging. Animals were injected with an intraperitoneal dose of 100 mg/kg D-luciferin (Promega, Charbonnières-les-Bains, France) and after five minutes, imaged with the IVIS Lumina S5 imaging system (Perkin Elmer, Villepinte, France). BLI data were analyzed using Living Image software (Perkin Elmer). BLI signal is reported as total flux in mouse (photons/sec). Throughout experiments, the body weight and score were monitored twice a week by operators blinded to treatment. Mice that survived past day 150 were considered long-term survivors. Once a week, blood samples were collected by retro-orbital sampling under local anesthesia. At the end of the experiment, mice were euthanized, tissue and blood samples were collected, and processed for immunophenotyping.

### Human primary cells immunophenotyping

For cellular immunophenotyping, T cells were stained with conjugated monoclonal antibodies (mAb) targeting HA, PD-1, TIGIT, TIM-3, CD271/NGFR (Miltenyi Biotec), LAG-3 and TGFBR2 (Biolegend, Amsterdam, Netherlands). For TILs characterization, expanded cells were stained with fluorophore-labeled antibodies against CD3, CD4, CD8, CD271/NGFR, PD-1, TIGIT, TIM-3, LAG-3, TGFBR2, CD45RA, CCR7, CD183/CXCR3, CD25, CD28, KLRG1, CD62L, CD127, FoxP3, CD40L, CD107a (Biolegend), TNFα (Miltenyi Biotec) and IFNγ (BD Biosciences, Heidelberg, Germany). After surface staining, FoxP3 was assessed after fixation and permeabilization using the BD Pharmingen™ Human FoxP3 Buffer Set (BD Biosciences). For determination of TILs functionality, expanded TILs were re-stimulated with dynabeads at a 1:3 bead:cell ratio in the presence of brefeldin A and the anti-CD107 staining antibody. After 18h, stimulation was stopped, and cells were stained after fixation and permeabilization with the eBioscience™ IC fixation buffer (Thermo Fisher Scientific, Hennigsdorf, Germany). For *in vivo* experiments, mouse spleen, and bone marrow samples were passed through a 70µm cell strainer to obtain a single cell suspension. Red blood cells were lysed with red blood cell lysis buffer (Merck Millipore, Fontenay-Sous-Bois, France). Next, cells were incubated with mouse Fc block (BD Biosciences) and stained with the conjugated antibodies targeting murine CD45, human TIGIT, human PD-1 (BD Biosciences), CD19-CAR FMC63, human CD19 and human CD3 (Miltenyi Biotec). Marker expressions were measured by flow cytometry and analyzed using FlowJo software (BD Biosciences).

### CAR-T cell activation, degranulation, and cytokine production assays

Transduced T cells were incubated with Nalm6-L-PD-L1 (ratio=1:3) in RPMI medium supplemented with 100U/ml of human interleukin-2 (hIL-2) at 37°C, 5% CO2. After 24 hrs, cells were harvested and stained with conjugated antibodies targeting HA, CD3, CD19, CD69, CD107a and PD-1 (Miltenyi Biotec). For intracellular cytokine quantification, 0.5µg/ml of Brefeldin A was added overnight to the coculture prior to downstream analysis. Cells were fixed, permeabilized with the FoxP3/Transcription factor staining buffer set (Thermo Fisher Sci.), and stained with conjugated antibodies targeting IL-2, IFNγ (BD Biosciences), TNFα, HA and CD3 (Miltenyi Biotec). Marker expressions were measured by flow cytometry and analyzed using FlowJo software (BD Biosciences).

### In vitro cytotoxicity assays

Cytotoxicity assays were performed by co-culturing CD19 CAR-T cells generated from three independent donors with the CD19-positive Nalm6-L-PD-L1 cell line at various effector to target (E:T) ratios. Briefly, 25,000 target cells were suspended in RPMI medium supplemented with hIL-2 (100U/ml) and CAR-T cells or PD-1 ZFR-CAR-T cells were added at the indicated E:T ratio (1:32, 1:16, 1:8 and 1:4). Untransduced T cells were used as control. After 24hrs at 37°C, 5% CO2, cells were harvested and living target cells were identified and quantified after staining with propidium iodide, anti-CD3 and negative for anti-CD19 (Miltenyi Biotec) to discriminate between effector and target cells. The specific lysis (%) was calculated as followed: 1-(N cancer cells at harvest / N cancer cells at seeding) x 100.

To assess anti-tumor cytotoxicity of transduced TILs, dNGFR-enriched TILs were co-cultured with SNU-C5-GFP-Luc cells at a 1:2 E:T ratio in DMEM supplemented with 1% FBS (Sigma Aldrich, Taufkirchen, Deutschland). Antigen-specific cancer cell killing by TILs was induced after addition of 1 ng/ml of a bispecific anti-CD3E-EpCAM antibody (ProteoGenix, Schiltigheim, France). After two days, cells were harvested, and viable SNU-C5-GFP-Luc cells were gently detached with TrypLE Express (Thermo Fisher Scientific) for 15 min. Harvested cells were stained 1:200 with Zombie UV™ Fixable Viability Kit (Biolegend) and anti-CD3-A700 (Biolegend) to distinguish TILs from GFP+ cancer cells. CD3E-EpCAM-mediated TILs cytotoxicity against SNU-C5-GFP-Luc cells was calculated based on cell counts of viable cancer cells relative to unstimulated cells co-cultured with TILs. TIL-specific killing was calculated as followed: 1 - (N viable cancer cells in presence of CD3E-EpCAM Ab / N viable cancer cells in absence of CD3E-EpCAM Ab) x 100.

### Statistics

Statistical analyses were performed using GraphPad Prism v.10.1.0 (GraphPad Software Inc., San Diego, CA, USA).

## Supporting information

David et al_ZFR_Onco_BioRxiv_Supplem

## DATA AVAILABILITY STATEMENT

All requests for data will be reviewed by Sangamo Therapeutics, Inc. to verify whether the request is subject to any intellectual property or confidentiality obligations. If deemed necessary, a material transfer agreement between the requestor and Sangamo Therapeutics, Inc. may be required for the sharing of some data. Any data that can be freely shared will be released.

## ACKNOWLEDGMENTS

The authors would like to thank the members of the animal care facility at Sangamo Ther. France (Valbonne, France), Jason Eshleman and Sarah Hinkley for the design and synthesis of ZFP encoding libraries at Sangamo Ther. (Richmond, CA, USA). We thank Duncan McKay, Jason Fontenot and Gregory Davis for strategic guidance and research organizational support. We also thank all patients for participation and all physicians from the department of Surgery, Campus Virchow-Klinikum, Charité-Universitätsmedizin, corporate member of Freie Universität Berlin, Humboldt-Universität zu Berlin for patient recruitment and providing CRLM tissues.

## AUTHOR CONTRIBUTIONS

M.D., P.S., D.M., G.S., A.E.M., C.F.D., C.J., I.M., S.K.T., J.V., L.N.T. and Y.Z. performed experiments. M.D., P.S., D.M., S.K.T., A.R., M.F. and D.F. designed experiments, analyzed and interpreted data. I.S. and W.S. provided tissue samples. G.D.D., I.K.N. and M.d.l.R. contributed to the discussions. MD, PS and DM contributed equally to the work and share first authorship, and the first-listed name was rotated across international meeting presentations and, accordingly, MD, PS, and DM agree and assert that any permutation of the order of these names is correct and acceptable. M.D., P.S., D.M., M.F., M.d.l.R. and D.F. wrote the manuscript. D.F. supervised the study.

