## Supplementary material for "Enhanced anti-tumor activity by Zinc Finger Repressor-driven epigenetic silencing of immune checkpoints and TGFBR2 in CAR-T cells and TILs": David et al_ZFR_Onco_BioRxiv_Supplem

### SUPPLEMENTAL MATERIAL AND METHODS

#### ZF-Repressor design and screening

ZFP backbones were designed using an archive of prevalidated one- and two-finger modules to produce a fusion with KRAB repressor domains as previously described[26]. ZFR mRNA transcripts were synthesized using the mMESSAGE mMACHINE T7 ULTRA Transcription Kit (Thermo Fisher Scientific). For the ZFR screening, human T cells were obtained from fresh Leukopaks, isolated using anti-CD4 and CD8 magnetic beads on a CliniMACS system (Miltenyi Biotec, San Jose, CA), aliquoted and stored in liquid nitrogen. After thawing, T cells were cultured in complete X-vivo15 (Lonza, Hayward, CA), supplemented with 100 IU/ml IL-2 (Thermo Fisher Scientific Waltham, MA), 5% Human AB Serum (Valley Biomedical, Winchester, VA), and activated with anti-CD3/CD28 Dynabeads (Thermo Fisher Scientific, Waltham, MA) at a bead:cell ratio of 1:3. T cell electroporation with 10µg ZF-Repressor mRNA was performed at Day 3 using a BTX ECM 830 Electroporator with Plate Handler HT-200 (BTX Harvard Apparatus, Holliston, MA). The transfection percentage was >90% as determined from co-transfection with a control GFP encoding mRNA. Two days after, ZFR-transfected T cells were lysed, mRNA content extracted and reverse transcribed to cDNA using a Power SYBR Green Cells-To-CT kit (Thermo Fisher Scientific) and qPCR reactions were performed using QuantiFast Multiplex PCR Master Mix (w/o ROX) (Qiagen, Redwood City, CA) as described previously[26]. The RT-qPCR probe/primer sets used are listed in Table S2.

#### Cell line cloning

Nalm6-L-PD-L1 cell line (See Figure S3) was obtained after transfection and puromycin selection of Nalm6-ffLuc (Creative Biogene, Shirley, NY, USA) with an in-house plasmid expressing PD-L1 and Puromycin resistance gene (pTXL-PD-L1-IRES-PuroR). Nalm6-L-PD-L1 cells were cultured in RPMI medium with 10% FBS + G418 (200 µg/ml).

SNU-C5 wild type cells were lentivirally transduced with green fluorescent protein (GFP)-firefly luciferase (ffLuc)_epHIV7 and followed by GFP sorting. Sorted SNU-C5-GFP-Luc cells were cultured in DMEM supplemented with 10% FBS and 1% Penicillin-Streptomycin (Merck, Darmstadt, Germany). Expression of EpCAM, PD-L1, CEACAM1, HLA-DR/DQ/DP (MHC-II), Galectin-3, CD112 and CD155 (Biolegend) was confirmed by flow cytometry and TGFβ secretion was detected using the ELISA kit U-PLEX TGFβ combo (human) kit (Meso Scale Diagnostics, Rockville, MA) according to manufacturer's instructions (See Table S3).

### SUPPLEMENTAL TABLES

**Table S1. CRLM Patient characteristics for TILs isolation**

| Pseudonym | Diagnosis | Gender | Age |
|---|---|---|---|
| mCRC01 | CRLM | f | 60 |
| mCRC02 | CRLM | f | 68 |
| mCRC03 | CRLM | m | 75 |
| mCRC04 | CRLM | f | 44 |
| mCRC05 | CRLM | m | 57 |
| mCRC06 | CRLM | f | 60 |
| mCRC07 | CRLM | f | 59 |
| mCRC08 | CRLM | m | 71 |
| mCRC09 | CRLM | f | 76 |
| mCRC10 | CRLM | f | 64 |
| mCRC11 | CRLM | m | 60 |
| mCRC12 | CRLM | m | 58 |
| mCRC13 | CRLM | m | 85 |
| mCRC14 | CRLM | m | 55 |

*Abbreviations: CRLM: colorectal cancer liver metastasis, f: female, m: male*

**Table S2. List of commercial assays used to monitor mRNA expression of the indicated genes.**

| **Gene** | **Assay ID** | **Dye** | **Supplier** |
|---|---|---|---|
| **Probes used to measure gene expression in mRNA electroporated T cells** | | | |
| *PD1* | Hs01550088_m1 | FAM | Thermo Fisher Sci. |
| *TGFBR2* | Hs00234253_m1 | FAM | |
| *TIGIT* | Hs00545087_m1 | FAM | |
| *TIM3* | Hs00262170_m1 | FAM | |
| *LAG3* | Hs00158563_m1 | FAM | |
| *EIF4A2* | Hs.PT.58.3053665 | HEX | Integrated DNA Technologies |
| *ATP5B* | Hs.PT.58.20452786 | Cy5 | |

**Table S3. Expression of cell surface markers and TGFβ family secretion in SNU-C5-GFP-Luc cell line.**

| | | Constitutive | | After 24h +100ng/ml IFNγ | |
|---|---|---|---|---|---|
| **IC** | **Marker / IC ligand** | **Positive cells [%]** | **MFI [sample-isotype]** | **Positive cells [%]** | **MFI [sample-isotype]** |
| - | GFP | 96.90 | | | - |
| - | EpCAM | 99.90 | 127899 | | |
| PD-1 | PD-L1 | 95.06 | 2988 | 98.41 | 7875 |
| TIM-3 | CEACAM1 | 45.04 | 729 | 64.61 | 1494 |
| LAG-3 | MHC-II | 6.93 | 48 | 12.03 | 133 |
| | Galectin-3 | 96.91 | 3644 | 94.88 | 2750 |
| TIGIT | CD112 | 99.64 | 8991 | 99.58 | 7857 |
| | CD155 | 99.96 | 37886 | 99.97 | 37544 |
| | **TGFβ1/2/3 ELISA** | **Concentration [ng/ml]** | | | |
| TGFBR2 | TGFβ1 | 2.96 ± 0.12 | | | |
| | TGFβ2 | 9.11 ± 2.18 | | | |
| | TGFβ3 | 0 | | | |

### SUPPLEMENTAL FIGURES

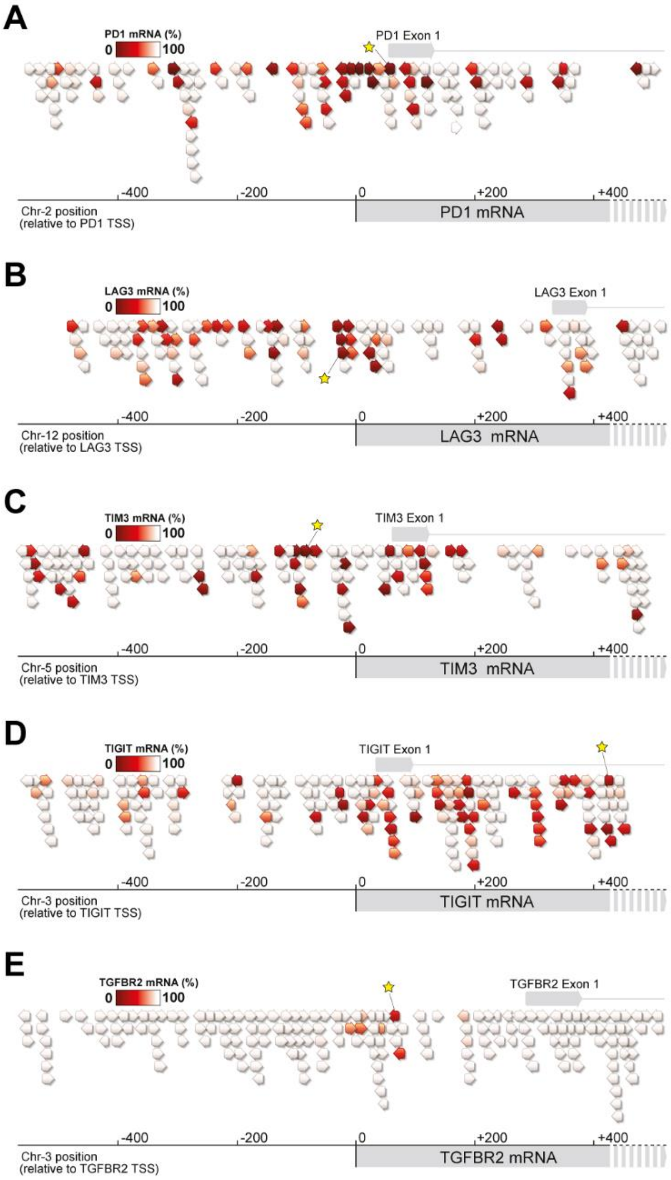

**Figure S1. Screening of functional ZF-Repressors in primary T cells targeting immune checkpoints and the TGFβ type 2 receptor.** Schematic of the *PD1*(A) *LAG3* (B), *TIM3* (C), *TIGIT* (D) and *TGFBR2* (E) target genes with ZFR candidates binding near the TSS. Arrows represent ZFR binding locations and orientations on the target gene and the red color intensity correlates with repression efficiency evaluated by RT-qPCR two days after ZFR mRNA transfection in T cells. The selected lead ZFRs are highlighted by a yellow star. Chr, chromosome; TSS, transcription start site.

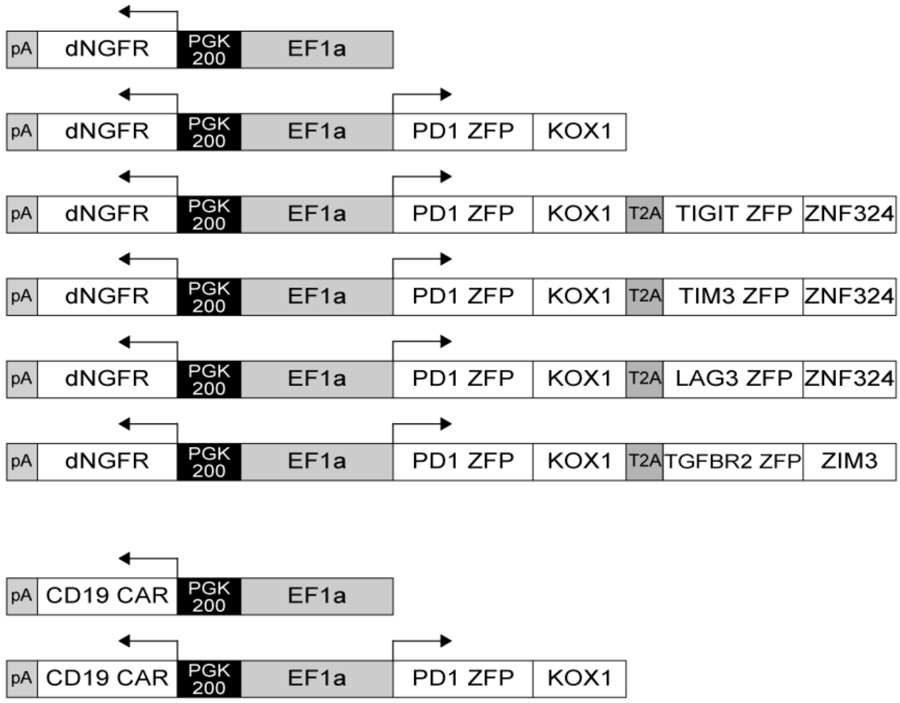

**Figure S2. Schematic representations of the lentiviral constructs used in this study**. pA, SV40 polyA; dNGFR, truncated human nerve growth factor receptor; human PGK200, short 200bp phosphoglycerate kinase promoter; EF1a, human elongation factor 1 alpha promoter; CD19 CAR, anti-human CD19 antigen. PD-1 zinc finger protein (ZFP) is to the KOX1 KRAB domain, TIGIT, TIM-3 and LAG-3 ZFP rea fused to ZNF324 KRAB domain, TGFBR2 ZFP is fused to ZIM3 KRAB domain.

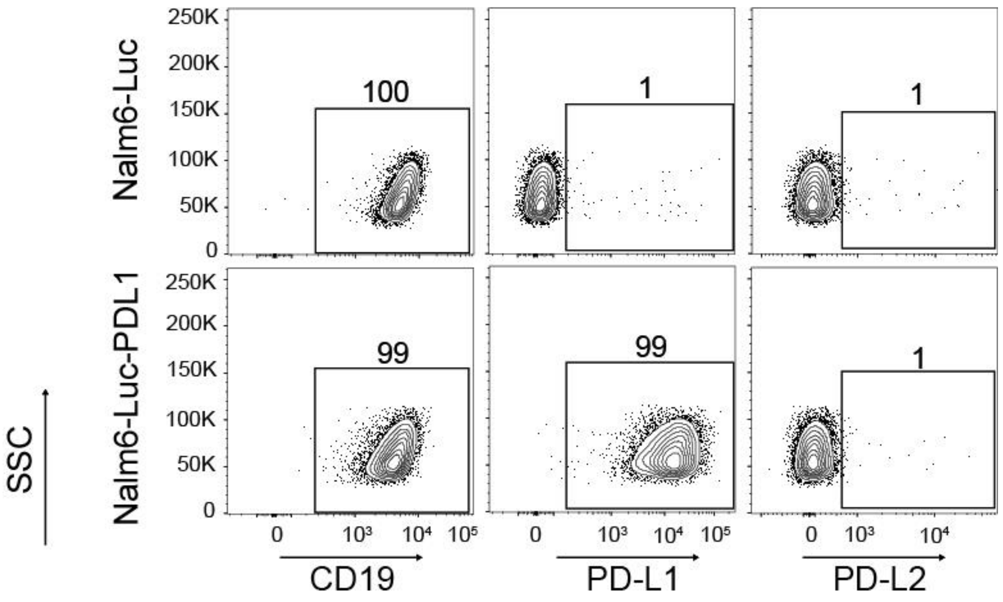

**Figure S3. Characterization of the Nalm6-L-PD-L1 cell line**

Cell surface expression of CD19, PD-L1, and PD-L2 markers in the parental Nalm6-Luc and the Nalm6-Luc-PD-L1 cell line was determined by flow cytometry.

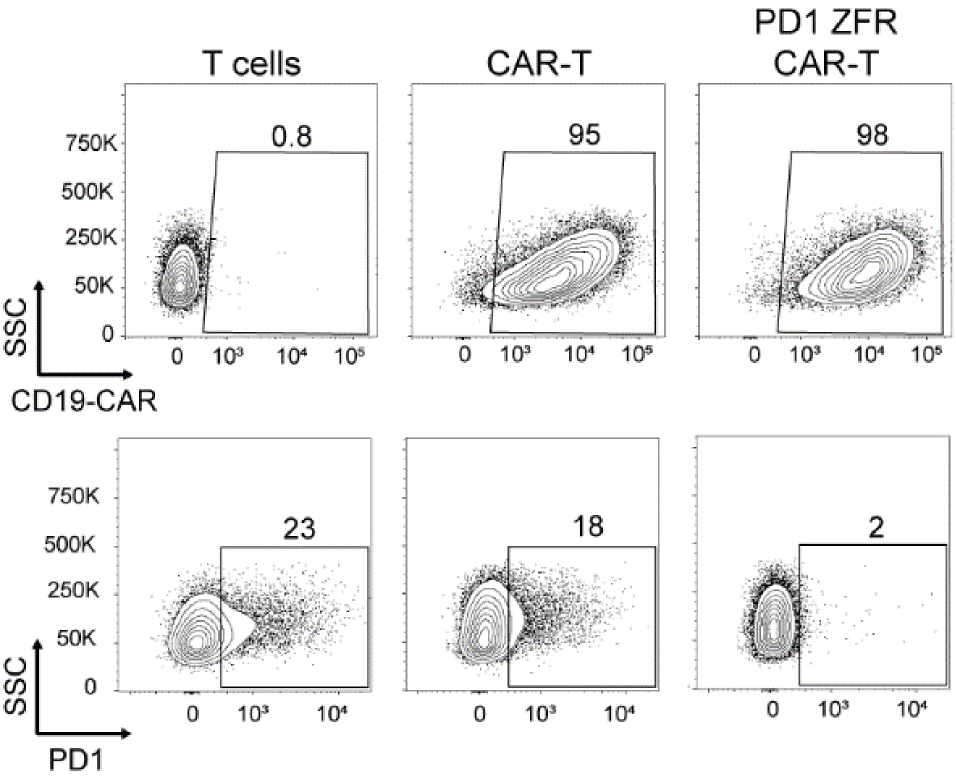

**Figure S4. Characterization of human ZFR-expressing T cells before *in vivo* injection.** Four days post-transduction with CAR or PD-1 ZFR CAR-expressing LV, CAR-positive T cells were enriched via magnetic cell sorting. Flow cytometry profiles are representative of CD19-CAR expression (top panel) and PD-1 expression levels (bottom panel) in CAR-T cells before *in vivo* injection at day 8.
